# Sparse Bursts Optimize Information Transmission in a Multiplexed Neural Code

**DOI:** 10.1101/143636

**Authors:** Richard Naud, Henning Sprekeler

## Abstract

Many cortical neurons combine the information ascending and descending the cortical hierarchy. In the classical view, this information is combined nonlinearly to give rise to a single firing rate output, which collapses all input streams into one. We propose that neurons can simultaneously represent multiple input streams by using a novel code that distinguishes single spikes and bursts at the level of a neural ensemble. Using computational simulations constrained by experimental data, we show that cortical neurons are well suited to generate such multiplexing. Interestingly, this neural code maximizes information for short and sparse bursts, a regime consistent with in vivo recordings. It also suggests specific connectivity patterns that allows to demultiplex this information. These connectivity patterns can be used by the nervous system to maintain optimal multiplexing. Contrary to firing rate coding, our findings indicate that a single neural ensemble can communicate multiple independent signals to different targets.

## Introduction

Visual, auditory and motor processing in the mammalian brain are organized in a hierarchy^1–5^. At the bottom of this hierarchy, ensembles of neurons code a dense array of simple features such as local visual contrast or simple movement components. At the top of the hierarchy neurons code more complex features such as complex images and movement sequences. Given that information travels both up and down the hierarchy with the power to drive or modulate responses^6–9^, we are compelled to an important question: How do populations that receive both bottom-up and top-down information process these two different types of messages?

Experimental observations argue for several opposing views. In one view, descending inputs modulate the bottom-up responses^7^. In a second view, top-down inputs can create responses *de novo*^8^. A third view arises from conceptual requirements. In the theories of unsupervised learning, the same units must simultaneously communicate feature recognition to higher-order units and a feature prediction to lower-order units^10,11^. In supervised learning, the higher-order success signal must percolate down the hierarchy, requiring units to communicate both the credit residual from top to bottom and an activation from bottom to top^12–14^. Also, in the binding problem, neurons are required to simultaneously signal the presence of a lower-order feature and its binding to a high-order one, across modalities^15–17^. Hence the third view is that of multiplexing: the same population needs to communicate different functions of ascending and descending information, simultaneously and to possibly different target neurons.

Present neural mechanisms for multiplexing can be separated in two different categories. First, spike-phase multiplexing^16,18^ posits that a population represents bottom-up information by its firing rate and top-down information by the timing of its spikes with respect to distinct frequency bands of a local field potential. This type of frequency-division multiplexing^19^ is supported by multiple experimental studies in different systems^16,18^, but the cellular mechanisms for encoding and decoding with the local field potential remain to be fully articulated. A second possibility is to allow the neurons to alternate between different modes: one devoted to the transmission of ascending information and another for the propagation of descending information. Time-division multiplexing of this kind is common in artificial neural networks^10,12,20^. In a similar fashion, time-division multiplexing is a useful mechanism in computational models of synaptic plasticity^14,20,21^, where the population alternates between sensing and learning phases. Yet it is not clear how time-division multiplexing can be mapped on the ongoing activity of cortical networks^9,20^.

In this article we propose a novel type of multiplexing based on the separation of bursts and single spikes at the level of an ensemble. Burst coding, we suggest, acts on the level of an ensemble to represent multiple information streams simultaneously and without ambiguity. We study this idea in the Thick-tufted Pyramidal Neurons (TPNs) as a paradigmatic cell type that receives both bottom-up and top-down signals. Using computational simulations, we show that TPNs can encode two independent streams of information with high temporal precision. The two streams can be decoded by post-synaptic populations using short-term plasticity and disynaptic inhibition. A theoretical analysis demonstrates that information representation is optimal for short and sparse bursts, a regime consistent with bursting in vivo. We further show that this optimal regime can be preserved by a network architecture that shares interesting parallels with the anatomy of dendritic feedback inhibition in the cortex. The proposed Burst Ensemble Multiplexing (BEM) code could allow to distinguish ascending from descending information and hence suggests a new approach to their analysis.

## Results

We consider a neural code where spike-timing patterns – single spikes and bursts – are separated at the level of individual spike trains before being averaged across a neural ensemble (Fig. 1a). From classical studies on the firing rate^22–24^, we expect that the resulting time-varying rates of single spikes and bursts can be related to the time averaged rates, but for time-varying stimuli they are generally not equal. How could rates of distinct spike-timing patterns represent different input streams or features? The simpler variant would be that single spikes and bursts are generated by two independent cellular mechanisms that each depend on one input stream alone. In this case, the ensemble singlet rate and ensemble burst rate would encode these streams independently. This possibility has been explored in the context of single cell firing of the thalamus^15,25^, hippocampus^26^, cortex^27,28^ and the electrosensory lateral lobe (ELL)^29–31^.

**Figure 1.**
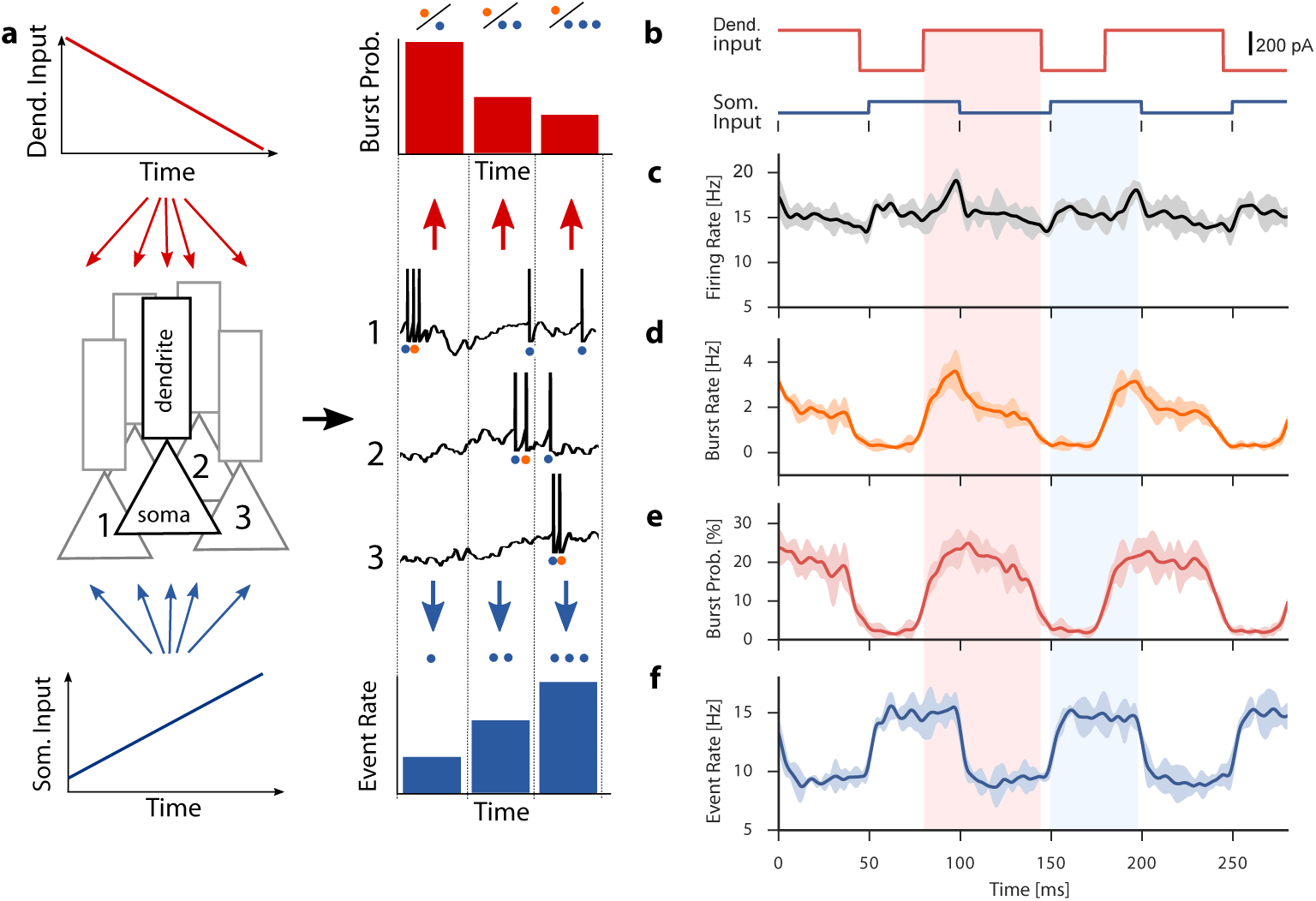
Burst ensemble multiplexing for representing simultaneously two signals. **a** Schema of the suggested neural code: One signal is delivered to the somata and another to the apical dendrites of a neural ensemble. Each neuron responds with a series of action potentials (black traces), which can be classified as isolated spike events or burst events on the basis of the interspike interval. The total number of events (blue dots) can be computed in each time bin to form an event rate (blue bars). The burst probability (red bars) is calculated by taking the ratio of the number of bursts (orange dots) and the total number of events for each time step. **b-f** Computational simulation: **b** Alternating somatic and dendritic input currents were used as inputs to a population of 4000 two-compartment neuron models. Phase lag and amplitudes were chosen such that **c** the output firing rate remains largely constant, to illustrate the ambiguity of the firing rate code. **d** The ensemble burst rate reflects a conjunction of somatic and dendritic inputs. **e** The burst probability reflects the alternating dendritic input. **f** The ensemble event rate reflects the alternating somatic input. Shaded regions show two standard deviations, calculated over five trials.

Alternatively, bursts could be generated by a synergy of the two input streams, namely *conjunctive bursting*. Cellular and molecular mechanisms for burst firing in the thalamus^15,32^, the superficial ELL^31^, L2-3 pyramidals^33^, CA1 pyramidals^34,35^, and TPNs^27^ can be said to burst in response to a conjunction of distinct streams of information. Since in this case both singlet rate and burst rate represent a mixture of the two input streams, contrasting singlet and burst rates is not likely to reveal independent information.

In TPNs, dendritic spikes convert a somatically induced singlet into a burst via the activation of a calcium spike in the dendrites^27^. Therefore, we reason that, in TPNs, a dendritic input stream is represented by the probability that a somatically induced spike is converted into a burst. On the ensemble level, this *burst probability* is reflected by the fraction of active cells that emit a burst (Fig. 1a). Then, a somatic input stream should be reflected in the rate of either singlet or burst events. We termed this quantity *event rate* (Fig. 1a) and it is calculated by taking the sum of the singlet rate and the burst rate. Importantly, this event rate equals the firing rate only in the absence of bursting, and is otherwise smaller.

Although burst coding was the focus of many theoretical^28,36–39^ and experimental^29–31,35,40,41^ studies and although ensemble burst coding may have been implied in some experimental studies^35,41^, its potential as a neural code for multiplexing has not been explored previously. In the following, we use computational modeling and theoretical analyses to show that the anatomy and the known physiology of the neocortical networks is consistent with this neural code for TPNs.

### Encoding: Dendritic Spikes for Multiplexing

To illustrate the BEM code in cortical ensembles, we first consider the firing statistics of model TPNs as they respond to alternating dendritic and somatic input shared among neurons (Fig. 1b). Individual TPNs are simulated using a two-compartment model that has been constrained by electrophysiological recordings to capture dendrite-dependent bursting (Fig. S1a-d^27^), a critical frequency for an after-spike depolarization (Fig. S1e-h^42^), and the spiking response of TPNs to complex stimuli *in vitro*^43^. In addition to the shared alternating signals, each cell in the population receives independent background noise to reproduce the high variability of recurrent excitatory networks balanced by inhibition, as well as low burst fraction and the typical membrane potential standard deviation observed in vivo^44–46^ (see *Materials and Methods*). As a result, simulated spike trains display singlets interweaved with short bursts of action potentials. Both types of events appear irregularly in time and are weakly correlated across the population (Fig. S2). In the example illustrated in Fig. 1, the dendritic and somatic inputs were chosen to yield an approximately constant firing rate (Fig. 1c). This illustrates the ambiguity of firing rate responses: the same response could have arisen from a constant somatic input. The burst rate (Fig. 1d) is also ambiguous as it signals the conjunction of somatic and dendritic inputs. However, this ambiguity can be resolved since a strong dendritic input is more likely to convert a single spike into a burst. Indeed, the event rate and burst probability qualitatively recover the switching pattern injected into the dendritic and somatic compartments, respectively (Fig. 1e-f). Thus it emerges that TPNs can simultaneously communicate many different functions of the somatic and dendritic inputs depending on the spike-timing patterns considered in the ensemble average.

To determine the dynamic range of this multiplexing, we characterize the input-output (I-O) function of the ensemble by simulating the population response to short 20-ms current pulses of varying amplitude, delivered simultaneously to all compartments (Fig. 2a-b). Consistent with earlier computational work and *in vitro* recordings of the time-averaged firing rate^28,39,47,48^, the ensemble firing rate grows nonlinearly with the somatic input, with a gain that is modulated by concomitant dendritic input (Fig. 2c). Also consistent with the single-cell notion that bursts signal a conjunction of dendritic and somatic inputs^27^, the ensemble burst rate in our simulations strongly depends on both somatic and dendritic inputs (Fig. 2d). The event rate increases nonlinearly with the somatic input (Fig. 2e), but is less modulated by the dendritic input than the firing rate, consistent with an encoding of the somatic input stream. The dynamic range of event rate is limited to small and moderate input strengths since the event rate saturates when somatic input is sufficiently strong to produce a burst in all the cells. Similarly, burst probability grows with the dendritic input strength with a weak modulation by the concomitant somatic input (Fig. 2f), consistent with our hypothesis that burst probability encodes the dendritic information stream. The dynamic range of burst probability is limited by small to moderate dendritic inputs since strong dendritic inputs produce bursts across the entire population, saturating the I-O function.

**Figure 2.**
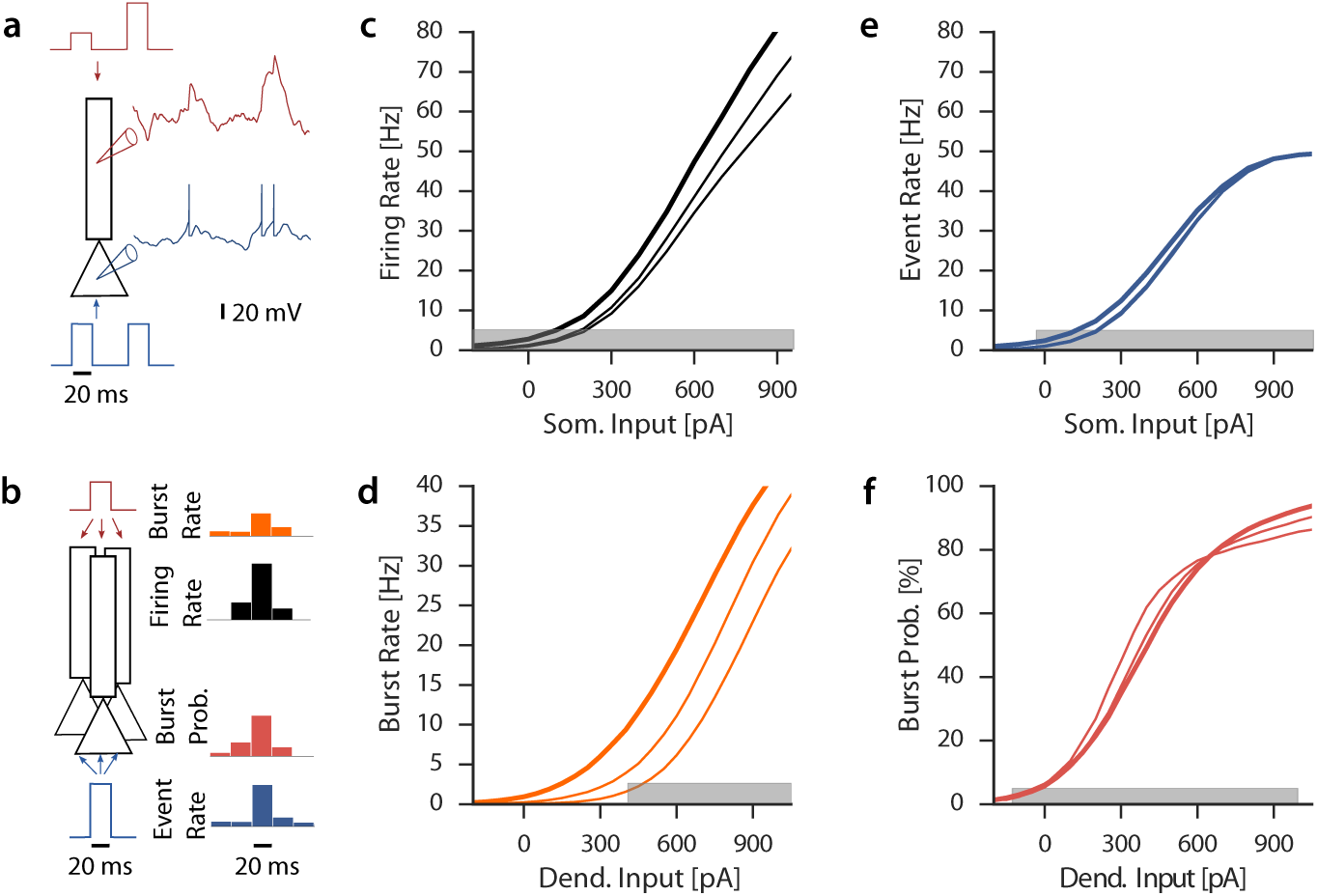
Input-output functions for distinct spike-timing patterns. **a** The two-compartment TPN model combines dendrite-dependent burst firing in the presence of high somatic and dendritic inputs with background noise. **b** Input-output functions are computed by simulating the response of 4000 TPNs to short current pulses and averaging across the ensemble. **c** The ensemble firing rate as a function of the somatic input amplitude is shown in the presence of a concomitant dendritic input (0, 200, 400 pA, thicker line corresponds to larger dendritic input). **d** The ensemble burst rate as a function of the dendritic input amplitude is shown in the presence of concomitant somatic input pulses (0, 200, 400 pA, thicker line corresponds to greater somatic input). **e** Same as **c** but for the ensemble event rate. **f** Same as **d** but for the burst probability, computed by dividing the ensemble burst rate by the event rate. Gray bars indicate the input regimes associated with SNR_*I*_>1 (see SI Data Analysis Methods).

To quantify the quality of this multiplexing, we compute a signal-to-noise ratio (SNR_*I*_), which is high for a response that is strongly modulated by the input in one compartment but invariant to input in the other (see SI Data Analysis Methods). We find that both the burst probability and the event rate reached larger SNR_*I*_ than either the burst rate or firing rate (maximum SNR_*I*_ > 250 for burst probability, and >1000 for the event rate vs SNR_*I*_ < 10 for the firing rate and <5 for the burst rate; Fig. S3). Also, the range of input amplitudes with an SNR_*I*_ > 1 is broader for burst probability than burst rate (gray regions in Fig. 2e-f). For very high inputs, the clear invariance of somatic and the dendritic input in event rate and burst probability breaks down (Fig. S4), because bursts can be triggered by somatic or dendritic input alone and are no longer a conjunctive signal (see Discussion). Therefore, multiplexing of dendritic and somatic streams is possible, unless either somatic or dendritic inputs are very strong. The low firing rates and sparse occurrence of bursts typically observed in vivo^44–46^ are in line with this regime, indicating that TPNs are well-poised to multiplex information.

### Information-Limiting Factors in Multiplexing

Given the need for fast cortical communication^49^, we ask if BEM is limited in terms of how fast it can encode two input streams. To this end, we simulated the response of an ensemble of TPNs receiving two independent input signals, one injected in the dendrites, the other injected in the somata. Both inputs are time-dependent and fluctuate with equal power in fast and slow frequencies over the 1-100 Hz range (SI Computational Methods). As a first step, we consider the case of a very large ensemble (80 000 cells) in order to minimize finite-size effects. Since the I-O functions obtained from pulse inputs (Fig. 2) are approximately exponential in the moderate input regime, we use the logarithm of the burst probability as an estimate of the dendritic input and the logarithm of the event rate as an estimate of the somatic input. Although crude compared to decoding methods taking into account pairwise correlation and adaptation^50–52^, this simple approach recovers accurately both the somatic and dendritic inputs (Fig. 3a-b), with deficits primarily for rapid dendritic input fluctuations. To quantify the encoding quality at different time scales, we calculate the frequency-resolved coherence between the inputs and their estimates. The coherence between dendritic input and its estimate based on the burst probability (Fig. 3c) is close to one for slow input fluctuations, but decreases to zero for rapid input fluctuations. Concurrently, the event rate can decode the somatic input with high accuracy for input frequencies up to 100 Hz (Fig. 3d). In both cases the coherence is at least as high, but typically surpasses the coherence obtained from the classical firing rate code, indicating that burst multiplexing matches but typically surpasses the information contained in the firing rate.

**Figure 3.**
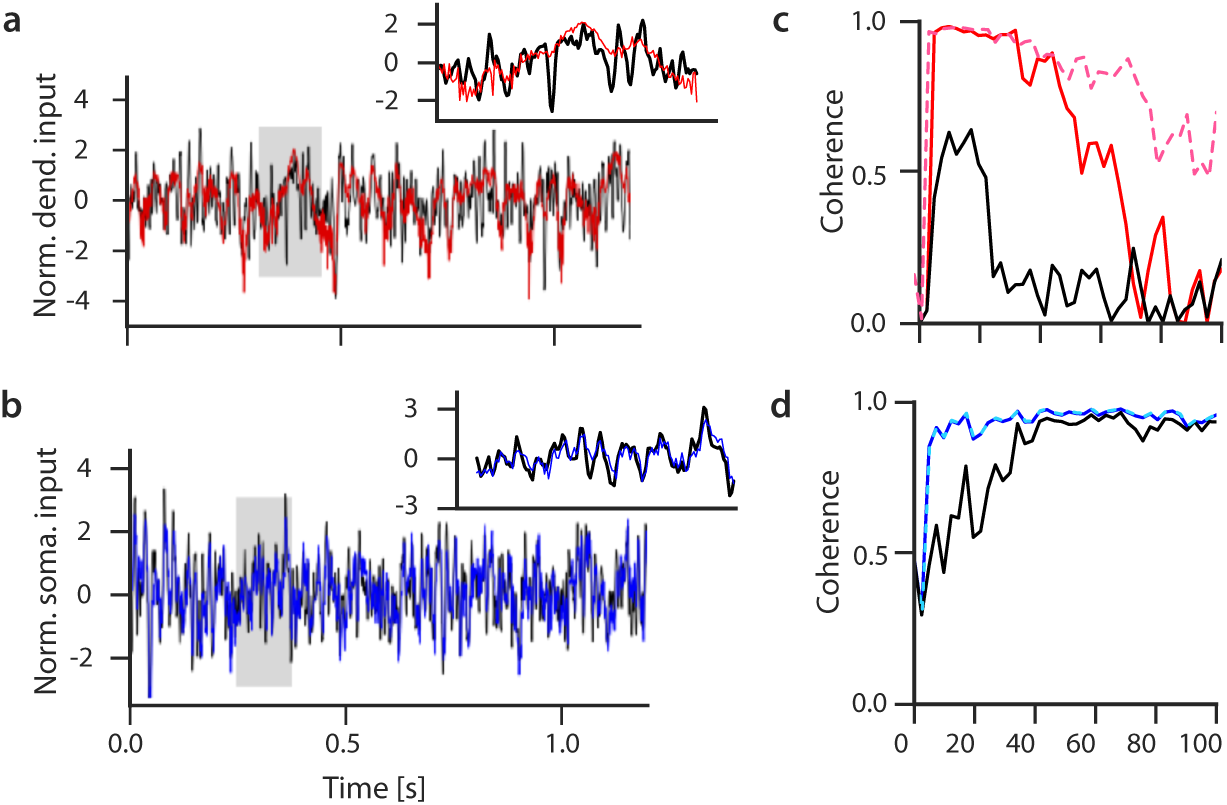
Decoding multiplexed time-dependent information. **a** The logarithm of the burst probability (red, normalized) is used as a decoded estimate of the dendritic input current (black, normalized). Inset: Blow-up of the shaded region. **b** The logarithm of the event rate (blue, normalized) is used as a decoded estimate of the somatic input current (black, normalized). **c** Frequency-resolved coherence between dendritic input and the logarithm of the burst probability (red) or the logarithm of the firing rate (black). The bandwidth of dendritic decoding increases for faster onset dynamics of dendritic spikes (red, dashed). **d** Coherence between somatic input and the logarithm of the event rate (blue) or the logarithm of the firing rate (black). Faster dynamics of dendritic spike does not affect the coherence with event rate (dashed cyan overlay). Decoding is performed on an ensemble of 80,000 cells, see SI Computational Methods for parameter values.

Previous studies that were not based on ensemble coding have shown that bursts encode the slowly varying part of sensory inputs only^31,53^. We therefore ask what limits the coherence bandwidth of the burst probability (Fig. 3c)? If it were limited by the dendritic membrane potential dynamics, coding could in principle be improved by changing membrane properties. If it were limited by the finite duration of bursts, which effectively introduces a long refractory period before the next burst can occur, this could introduce a fundamental speed limit for BEM. To investigate the latter, we performed an information-theoretic analysis of the BEM code, which indicates that the refractory period does not affect the bandwidth of BEM for sufficiently large ensembles, consistent with previous theoretical work^54^.

Alternatively, the membrane dynamics in the dendrite could limit the bandwidth. A slow passive dynamics in the apical dendrites is not likely to be a limiting factor, because the high density of the hyperpolarization-activated ion channels^55^ contributes to a particularly low dendritic membrane time constant^43^. The other possibility is that the slow onset of dendritic spikes limits the bandwidth^56^. This possibility would be compatible with slow calcium spike onsets observed, arising from the kinetics of calcium ion channels^57^. Therefore, we simulated the response to the same time-dependent input shown in Fig. 3 but with a three-fold increase of the voltage sensitivity for dendritic spikes, to accelerate the onset of dendritic spikes. This single manipulation considerably improved the encoding of high-frequency fluctuations (Fig. 3c-d and Fig. S6). We conclude that one important temporal limitation to burst coding in TPNs is the slow onset of dendritic spikes.

The total amount of information transmitted depends on a variety of extrinsic and intrinsic factors. The main extrinsic factors are the power and the bandwidth of the input signals. High power is manifested in large input changes, which can strongly synchronize cells, and can thus increase the ensemble response. Consistently, the information rates obtained in the previous section depend strongly on our choice of input power. For instance, decreasing the relative power in the dendrites decreases the information rate obtained from the burst probability (Fig. S5). To arrive at a more objective assessment of multiplexing, we derived mathematical expressions for the event and burst information rates at matched input power and total number of spikes (SI Theoretical Methods). It allows us to determine how coding depends on the properties and prevalence of bursts as well as the conditions under which multiplexing is advantageous.

Firstly, we find that there is an optimal burstiness, i.e., mean burst probability, for which information transmission is maximized (Fig. 4a). This optimum arises from the fact that rare bursting sacrifices information from the dendritic stream, whereas frequent bursting must sacrifice information from the somatic stream to meet the constraint of total number of spikes. This optimal burstiness depends on the number of spikes in a burst and the bandwidth of the two channels. It decreases with the number of spikes per burst (Fig. 4b), in line with the notion that long bursts convey little information per spike and should hence be used more sparsely. The optimal burstiness further decreases with decreasing dendritic bandwidth (Fig. 4b), that is for neurons with slower dendritic dynamics. The information transmitted decreases with the number of spikes per burst (Fig. 4c), in line with the intuition that the second spike in a burst marks the event as a burst, whereas additional spikes contain no further information. Hence, for neurons with slow dendritic dynamics, BEM performs best when bursts are short and occur rarely, in line with experimental observations^44^. Finally, the preference for short bursts is independent of the number of neurons in the ensemble (Fig. 4c), but a minimal number of neurons is required to transmit more information than a rate code with the same firing rate and number of neurons (Fig. 4c-d). If the somatic and dendritic compartments have the same bandwidth, the total information transmitted by BEM approaches twice the information of a classical rate code, in the limit of very large ensembles (Fig. 4d). In summary, the theoretical analysis suggests that short and sparse bursts in a large ensemble maximize information transmission in burst multiplexing.

**Figure 4.**
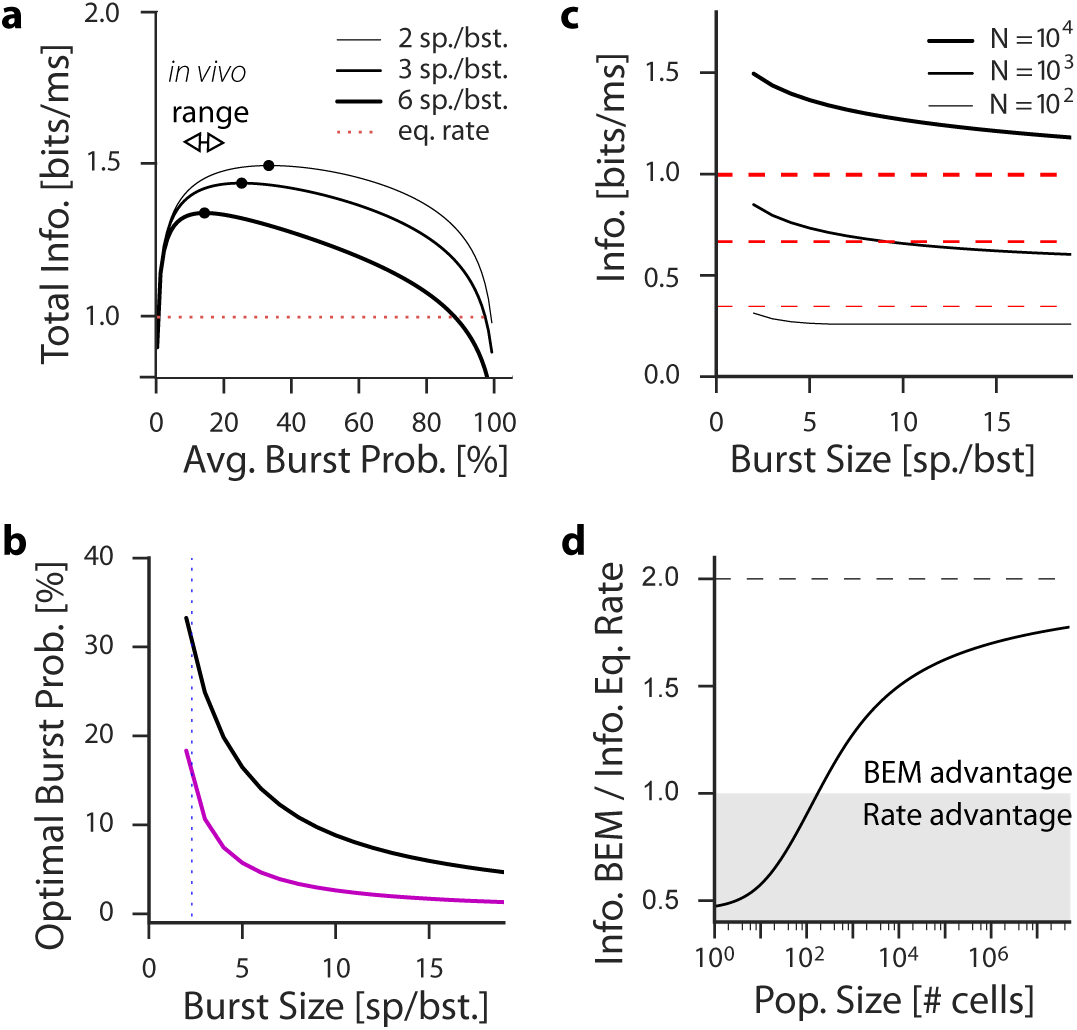
Short and sparse bursts are optimal for multiplexing. **a** Theoretical estimates of total multiplexed information constrained to a fixed total number of spikes (black curves, Eqs. S17, S29 and S30). The information varies as a function of the stationary burst probability and has a maximum for low burst probability (black circles), consistent with bursting statistics *in vivo*^44^. The three black lines correspond to three burst sizes (2, 3 and 6 spikes per burst), illustrating that the smallest burst size communicates the greatest amount of information. The information rate for a firing rate code with matched input amplitude and stationary firing rate is shown for comparison (red dashed line, Eq. S12). Parameter values were *p* = 0.5, *P* = 1, *N* = 10^4^, *W*_*s*_ = *W*_*d*_ = 100 Hz and *A*_0_ = 10 Hz. **b** The optimal mean burst probability decreases as a function of burst size. It reaches 31% for typical burst size (corresponding to an average of 2.3 spikes per burst^44^, dotted line) for parameters as in panel a. Considering a slower dendritic dynamics *W*_*d*_ = 0.2*W*_*s*_ reduces the optimal burst probability (magenta). **c** The maximum information rate (solid black curves, N=10^2^, 10^3^ and 10^4^) decreases as a function of burst size. It surpasses the information of the firing rate code with matched input amplitude and stationary firing rate (red lines, for corresponding *N*) for sufficiently large ensembles and for small burst sizes. **d** For matched input amplitude and output rate, the total multiplexed information gain relative to the information rate of the firing rate (Eq. S30 over Eq. S12) asymptotes to two (dashed line). The area where the classical firing rate offers an advantageous coding strategy is shown with shaded gray.

### Decoding: Cortical Microcircuits for Demultiplexing

For the brain to make use of a multiplexed code, the different streams have to be decodable by biophysical mechanisms. Previous experimental^58,59^ and theoretical^53,60^ studies have argued that short-term plasticity (STP) can play a role in facilitating or depressing the post-synaptic response of bursts. We have argued that ensembles of TPN may represent distinct information not in the rate of single spikes and bursts, but in the event rate and the burst probability. To test if STP can be used to recover these ensemble features, we simulated cell populations receiving excitatory input from TPNs, and studied how the response of these post-synaptic cells depends on the input to TPNs. As a model for STP, we used the extended Tsodyks-Markram model^59^ with parameters constrained by the reported properties of neocortical connections^61^ (see SI Computational Methods). By decreasing the postsynaptic effect of additional spikes in a burst, short-term depression (STD) could introduce a selectivity to events, particularly for short bursts and strong depression. Indeed, we find that in a population of cells receiving excitatory TPN inputs, responses correlated slightly more with event rate when STD was present than without STP (Fig. S7a-b). The presence of STD affects the range of SNR_*I*_ above one only weakly, but increases the maximum reached by SNR_*I*_ considerably (SNR_*I*_ reaches 120 with STD, and is below 30 without STD, Fig. S8a-b). STD can hence be interpreted as an event rate decoder. In turn, since TPN event rate encodes the somatic stream (Fig. 2), it is not surprising that we find STD to further suppress the weak dependence on dendritic inputs, while maintaining the selectivity to somatic input (Fig. 5f, h). Hence STD improves the selective decoding of the somatic stream.

**Figure 5.**
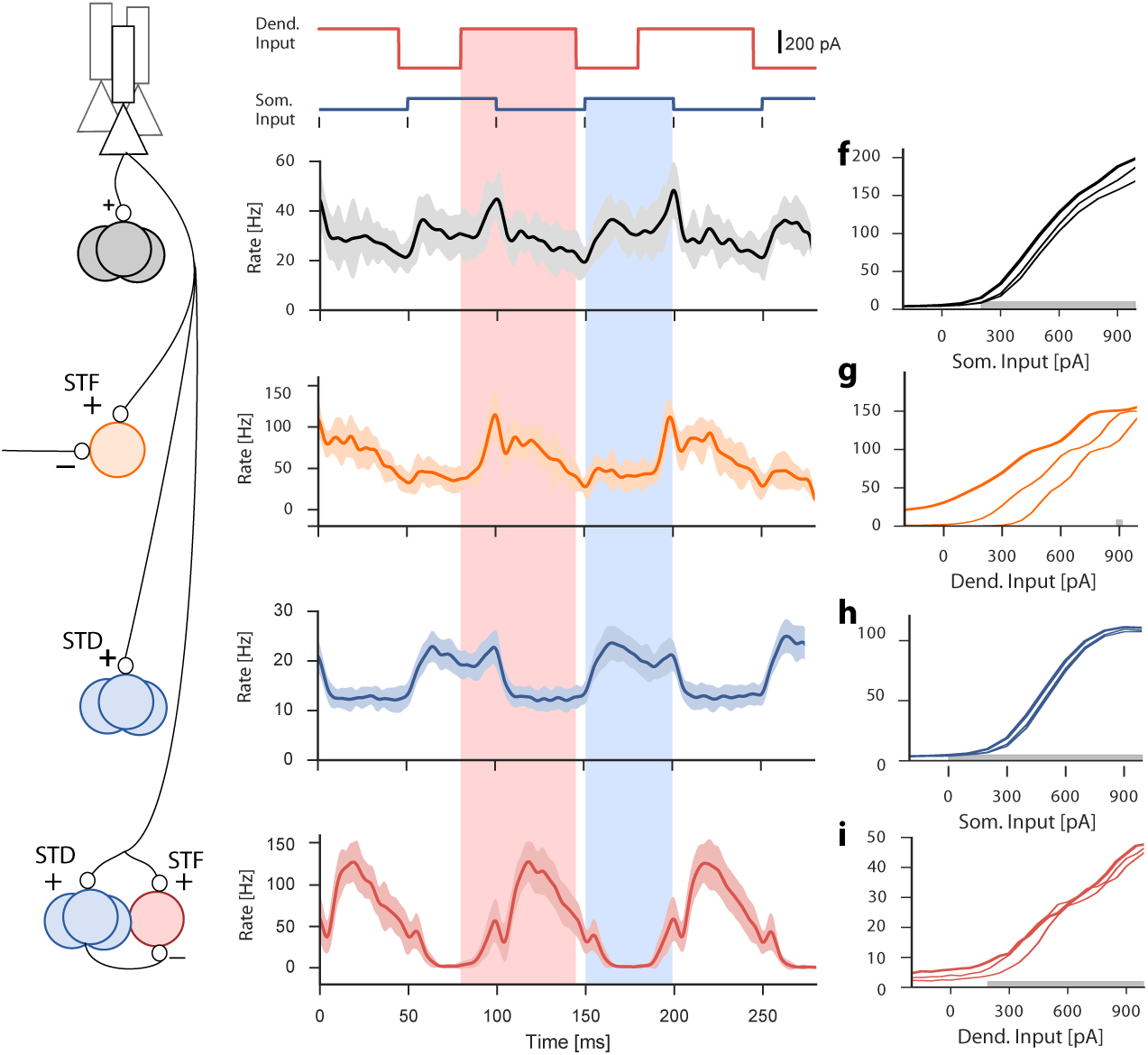
The role of short-term plasticity and disynaptic inhibition for separating distinct information streams. **a** Alternating somatic and dendritic inputs are injected in 4000 TPNs (as in Figure 1). The firing rate response of neurons post-synaptic to TPNs is shown for synapses **b** without STP, **c** with STF and constant inhibition, **d** with STD, **e** with STF and disynaptic inhibition. The responses of the post-synaptic cells are shown as a function of the amplitude of somatic and dendritic pulses. Firing rate of population receiving TPN input **f** without STP, **g** with STF and constant inhibition, **h** with STD, **i** with STF and disynaptic inhibition. The different lines show the response in the presence of a 200, 300 and 400 pA concomitant input (somatic input when abscissa is dendritic, and vice versa). Shaded area in in **c**-**g** shows two standard deviations around the mean. Thicker lines in **f**-**i** corresponds to the strongest concomitant stimulation. Gray shade in **f**-**i** highlights range with SNR_*I*_ > 1.

We then ask if post-synaptic neurons can decode the conjunction of inputs and the dendritic input streams. By increasing the postsynaptic effect of later spikes in a burst^58,60,62^, short-term facilitation (STF) boosts the sensitivity to bursts and hence to the dendritic stream (Fig. 5c,g). To decode the dendritic stream, neurons should compute a quantity similar to the burst probability (Fig. 2d). We reason that neural computation of burst probability could be achieved by combining burst rate sensitivity with divisive disynaptic inhibition from an event rate decoder. Thus, we consider a population receiving facilitating excitatory input from TPNs, combined with disynaptic inhibition from an STD-based event rate decoder. We manually adjusted the weights of these connections to increase the post-synaptic effect of dendritic inputs, while decreasing the post-synaptic effect of somatic input. This was achieved with potent excitation and inhibition, a regime associated with divisive inhibition^36,63^. The output rate of this microcircuit displayed a higher correlation with burst probability than for a microcircuit without disynaptic inhibition (Fig. S7c-d). This microcircuit can selectively decode dendritic input (Fig. 5i) with an SNR_*I*_ above 1 over a large range of dendritic input amplitudes (Fig. S8d). In the absence of disynaptic inhibition, the SNR_*I*_ reached one for a very small range of dendritic input amplitudes (Fig. S8c). Is the presence of STD essential to this operation? In line with the weak dependence of TPN firing rate on dendritic input, we find that the presence of STD in this microcircuit is not essential since decoding of the dendritic stream can also result from STF combined with disynaptic inhibition without STD (Fig. S8e-f). We conclude that a microcircuit with STP and disynaptic inhibition in a divisive regime can selectively extract different input streams from a multiplexed neural code.

### Gain Control of Multiplexed Signals

To transmit significant information, the burst code relies on a graded increase of the burst rate as a function of dendritic and somatic inputs, which is at odds with the all-or-none nature of calcium spikes in single cells^27^ and ensembles^28^. Three mechanisms can linearize the input-output function and transform an all-or-none response into a graded one. These mechanisms are background noise, spike-frequency adaptation and feedback inhibition. For fast and reliable encoding, feedback inhibition is the most efficient since linearization is faster than with adaptation and the signal-to-noise ratio better than with background noise. Feedback Dendritic Inhibition (FDI) is mediated in the neocortex by somatostatin-positive neurons (SOMs), which receive input from TPNs^62^ and project back to the apical dendrites of the same ensemble. FDI is known to linearize dendritic activity^64,65^ and may therefore linearize the burst probability, while feedback somatic inhibition may linearize the event rate. But since SOMs are activated by the TPNs, both somatic and dendritic inputs may reduce the burst probability, breaking the segregation of dendritic and somatic streams. Therefore, we ask whether FDI inhibition can linearize the burst response without introducing a coupling between the two input streams.

To this end, we simulated TPNs receiving feedback inhibition from a burst-probability decoder (Fig. 6a). We find that the presence of such FDI reduces the average burst length (Fig. 6b), consistent with similar experimental manipulations in the hippocampus^66^. Also, there is a weaker gain modulation of the firing rate I-O function compared to TPNs without FDI (Fig. 6c). Inhibition from the burst probability decoder motif does not abolish bursting in the TPNs but reduces both the overall proportion of bursts and the gain of the burst probability I-O function (Fig. 6d). Importantly, this form of FDI does not change the invariance of the burst probability to somatic input, so the multiplexed code is conserved (Fig. 6d). We suggest that this invariance requires that FDI is primarily driven by dendritic input to TPNs. The effect of somatic input to TPNs is counteracted by the disynaptic inhibition. When divisive disynaptic inhibition in the burst-probability decoder is replaced by a constant hyperpolarizing current (Fig. 6e), FDI becomes strongly modulated by somatic input (Fig. S9h). Although bursts remain shorter (Fig. 6f) and sparser (Fig 6h), and although the gain of the firing rate response is reduced in a very similar fashion (Fig. 6g), the burst probability loses its invariance with respect to somatic input (Fig. 6h). Therefore, FDI from the burst probability decoder motif contols the gain of the dendritic signal and ensures that bursts are short and sparse, while maintaining the suggested multiplexed code.

**Figure 6.**
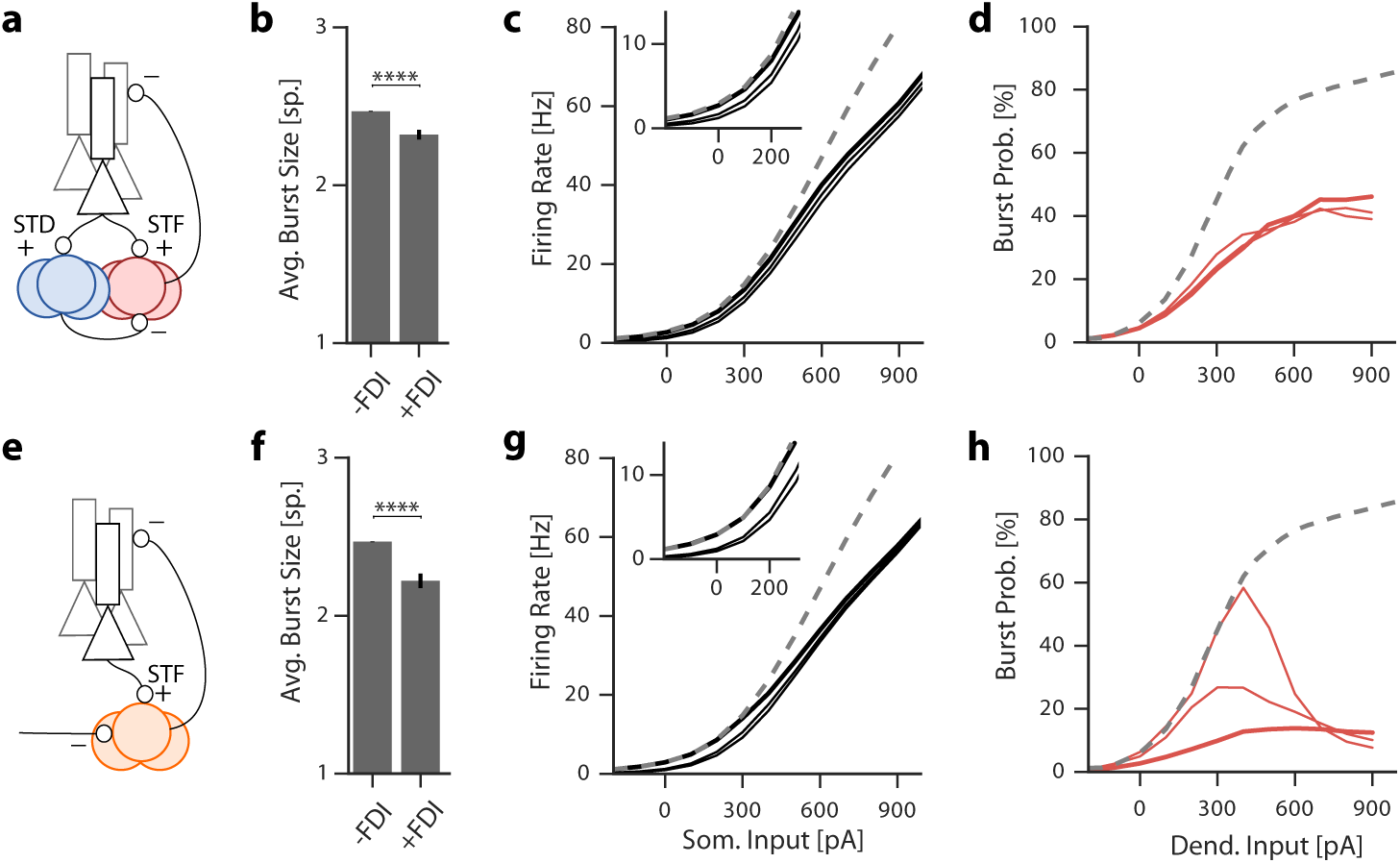
Dendritic feedback inhibition controls multiplexing gain. **a** Schematic representation of the simulated network. **b** Burst size averaged across simulated input conditions with and without feedback dendritic inhibition (FDI). **c** Dependence of TPN firing rate on somatic input for three different amplitudes of dendritic input pulses (0, 200 and 400 pA), inset shows zoom close to firing rate onset. **d** Dependence of burst probability on the dendritic input for three different somatic input amplitudes (0, 200 and 400 pA). The thicker line shows the response with the strongest somatic input. **e**-**h** As above, but replacing disynaptic inhibition by a hyperpolarizing current (430 pA). In this case, the burst probability associated with small dendritic inputs is decreased by somatic input (**h**). Stars indicate significant difference (Welch’s t-test p <0.0001). Dashed line shows response in the absence of FDI (corresponding to thick lines in Fig. 2).

## Discussion

We have introduced a neural code able to simultaneously communicate two streams of information through a single neural ensemble. This neural implementation of multiplexing is distinct from time-division^20,67^ and frequency-division multiplexing^19^ and is specific to communication with spike trains. Contrary to single-cell burst coding^31^, we have found that ensemble burst coding can encode quickly changing inputs, although processing speed may be limited by the biophysical properties of active dendrites. This code is optimal for short and sparse bursts, which is consistent with observed bursting in L2-3 and L5B cells^44^. Finally, we have illustrated in simulations how ensemble multiplexing suggests specific connectivity motifs to demultiplex burst coded information. We believe that this neural code satisfies the need to communicate different quantities in top-down and bottom-up direction through the same neurons.

## Extensions of Multiplexing

Is BEM antagonizing frequency-division multiplexing? Frequency-division multiplexing has been suggested on the basis of experimental observations^16,18^. Burst coding can in theory supplement this type of code since distinct types of event may synchronize to distinct frequency bands. Since this idea is supported by experiments^68^, we believe BEM does not exclude additional multiplexing using frequency-division of the local field potential.

We have focused on the information content of event rate and burst probability because these features of the ensemble response can be controlled independently. The nervous system can use this multiplexing to send to different targets, different functions of the bottom-up and top-down inputs. As we have shown, it can be achieved by tuning the properties of STP as well as the synaptic weights connecting inhibitory cells. This opens up a large range of possible computations. In one extreme example, top-down and bottom-up information can be communicated through the same neurons without affecting each other, as described schematically in Fig. 7a. In another example, the nervous system may have to modify descending information as a function of bottom-up information, as illustrated schematically in Fig. 7b. We note that both examples are consistent with the theory that top-down input modulates the firing rate response^7^, since the firing rate is modulated by top-down input to the dendrites (Fig. 2c). Furthermore, when calcium indicators are used to report activity levels, both examples can also be consistent with the theory that top-down input drives responses, since calcium indicator may report the burst rate^69^. In order to resolve the computation performed by descending inputs, it appears essential to distinguish different spike timing patterns.

**Figure 7.**
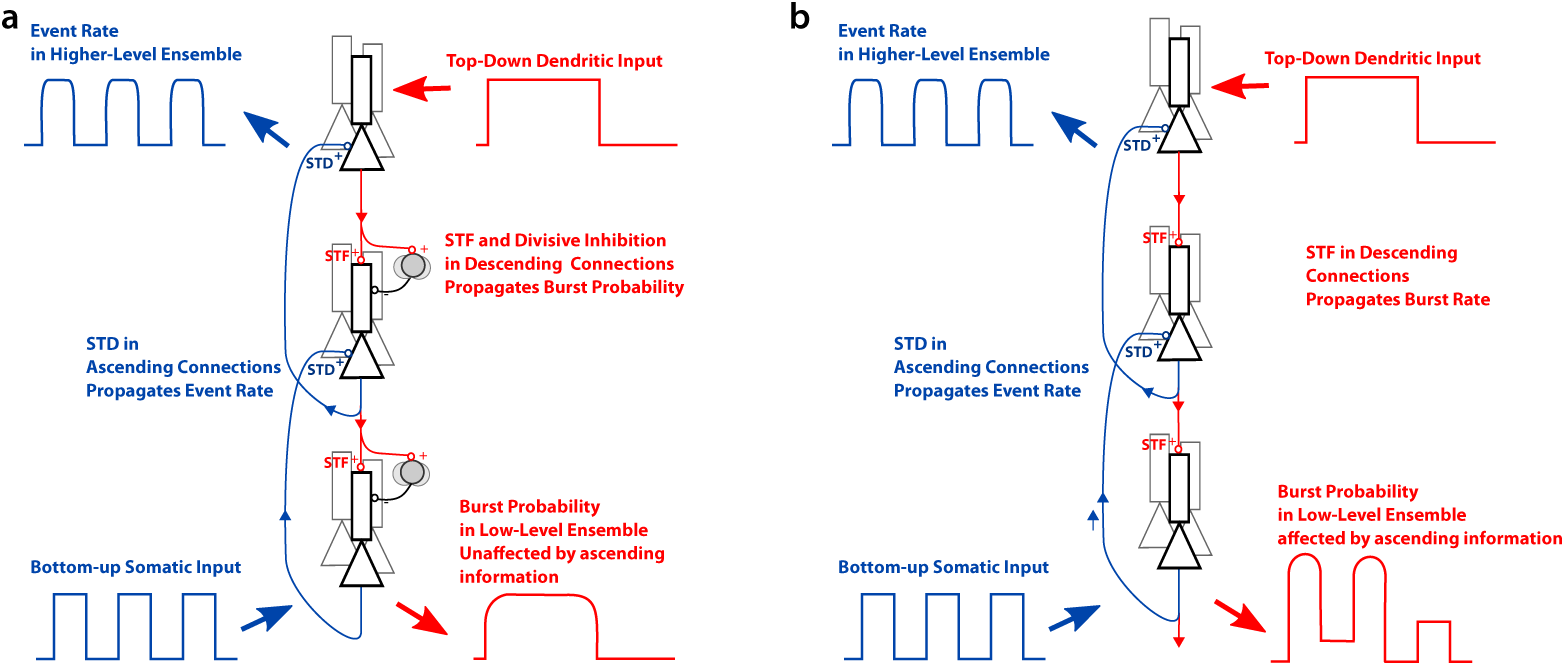
Potential functional roles of burst ensemble multiplexing. **a** When the descending connections have STF with disynaptic inhibition, top-down information can propagate down unaltered by bottom-up information, the burst probability. Ascending connections with potent STD can communicate bottom-up information even in the presence of potent descending drive. **b** When descending connections have STF without potent disynaptic inhibition, a conjunction of ascending and descending information, the burst rate, is communicated down.

For clarity of the exposition, we have limited our discussion to a distinction between singlets and bursts. It is has been suggested that the size of bursts may encode additional information^26,70^. One can generalize BEM to capture three distinct inputs controlling three types of events, namely singlets, short bursts and long bursts. There is a parallel between these three types of events and the three types of local regenerative activity in TPNs^71^. We can speculate that encoding three streams of information would be possible when calcium spikes generate short bursts and when these short bursts can be converted into long bursts by slower NMDA spikes. A postsynaptic decoding of such a complex temporal ensemble code would probably require intricate dynamics of STP, however, potentially combined with micro-circuits.

### Limitations of Multiplexing

The multiplexing mode of burst coding described here is limited to a regime where the inputs to be multiplexed are small or moderate. In TPNs, strong somatic inputs could trigger bursts of action potentials, which would challenge an association between dendritic inputs and burst probability. Since the regime where multiplexing can arise (see Fig. 2) is clearly reflected in the ensemble event rate and burst probability, experiments could assess whether TPNs are maintained in this regime or if the ensemble can switch between multiplexing and classical rate coding. Whether the inhibitory motifs described here can pick up this switch of modes is a question that lies beyond the scope of this study.

Another limitation lies in the assumption that spikes are probabilistically converted into bursts independently across neurons. Dendrites should hence be in an asynchronous state with weak pairwise correlations. This asynchronous state was suggested to enable rapid^72^ and efficient^73^ encoding. In our simulations, the asynchronous state was mimicked by a substantial background noise that effectively desynchronized and linearized the responses of both somatic and dendritic compartments. A physiological basis for this background noise could lie in the stochastic activation of ion channels^74–76^ as well as in a state of balanced excitation and inhibition, which is known to favor pairwise decorrelation^45,77^ and could be supported by homeostatic inhibitory synaptic plasticity^78^. We have argued that FDI can keep TPNs in a state of short and sparse bursts, it is natural to suggest that FDI can also ensure asynchronous dendritic activity. Burst ensemble coding hence requires a synergistic coordination among single cell bursting mechanisms, the morphological targets of different input streams on these cells, and neuronal circuit motifs. Whether hallmarks of such a coordination can be found in different neuronal systems, how this synergy is established and maintained by plasticity and modified by neuromodulation remains an open question.

### Neural Implementations of Multiplexing

We suggested that a BEM code can be decoded by microcircuits combining STP and inhibition. Candidates to implement this microcircuit are local GABAergic cells. These cells receive input from local TPNs and are known to interconnect into specific microcircuits. Consistent with the circuit motifs described in Fig. 5, cortical cells typically receive facilitating inputs from TPNs^59,79,80^, while parvalbumin-positive (PV) cells and vaso-intestinal peptide-positive (VIP) cells typically receive depressing TPN input^59,79,80^. Both PV or VIP cells can therefore encode event rate, but given that VIP cells share direct top-down inputs with TPN apical dendirtes^81,82^, it is more likely that PV cells relate to the somatic input.

Compatible with the burst probability decoder in Fig. 5e, disynaptic inhibition onto SOM cells arises from either PV or VIP cells^83,84^. A sub-type of SOM cells, the Martinotti cells, inhibits specifically the dendrites^85^ and acts as a powerful control of dendritic spikes^27,64,65^, consistent with the motif used in Fig. 6a. This suggests that Martinotti cells are well-poised to encode burst probability. Our theoretical analysis suggests that the burst code is limited to slower signals and requires larger ensembles. This is consistent with the anatomy; Martinotti cells have slower dynamics and project back to larger ensembles. Both these predictions are verified by the anatomy^86,87^ and the electrophysiology^62,80^ of SOMs. Another testable prediction in this context is that Martinotti cells must receive disynaptic inhibition to preserve multiplexing of the local ensemble (see Fig. 6), this property may arise concurrently with calcium spikes in development.

Connection motifs that could decode multiplexed codes are not restricted to neurons in the vicinity of TPNs. We hypothesize that such circuits are beneficial wherever TPN outputs need to be interpreted. For example, long-range connections from the cortex to the thalamus show a combination of STF and disynaptic inhibition^88^.The proposed relationship between TPN bursting and connectivity can be tested in the remote targets of TPNs^89^.

Our work was motivated by TPNs in layer 5B of the somatosensory cortex, which receive distinct input streams on their somatic and dendritic compartment^90,91^, have ascending and descending projections^6^ and show dendrite-dependent bursting^27^. Our work suggests that the input onto these apical dendrites (e.g, by attentional signals^68,92^ or other top-down signals^6,90^) should be represented in the ensemble burst probability. Consistent with this prediction, burst fraction in somatosensory cortex correlates with the ability of mice to report a light touch^93^. In higher-order cortex, elevated burst fraction is associated with attention in primate frontal cortex^68^, which offers an interesting parallel. Also in the electrosensory lobe of the electric fish, top-down input enhance burst generation^41,94,95^.

### Conclusion

Cracking the cortical neural code is to attribute proper meaning to temporal sequences of action potentials. Important clues to this riddle are the biophysical mechanisms that mediate the encoding and decoding of information in spike trains. We have linked here features of the physiology of the cortex with the optimization of a multiplexing neural code. The code described here suggests that the opposing views of top-down input as either modulating^7^ or driving responses^8^ can be reconciled by a multiplexed neural code.

## Materials and Methods

Bursts are defined as a set of spikes followed or preceded by an interspike interval smaller than 16 ms. Burst rate is calculated by finding all bursts and summing across the population. Event rate is calculated by finding all isolated spikes and the first spike in a burst before summing across the population. Burst probability is calculated as the ratio of the burst rate over the event rate. A smoothing kernel of 10 ms is used for displaying the time-dependent rates.

### Network simulations

The network consists of four types of units: pyramidal-cell basal bodies, pyramidal-cell distal dendrites and inhibitory cells from population a receiving STF and b receiving STD. We used the lower case letters s, d, a and b to label the different units, respectively. Somatic dynamics follow generalized integrate-and-fire dynamics described by a membrane potential *u* and a generic recovery variable *w* to account for subthreshold-activated ion channels and spike-triggered adaptation^24^. For the ith unit in population *x* we used
(1)

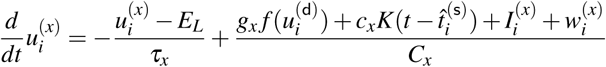

(2)

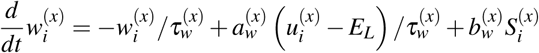

where *E*_*L*_ is the reversal potential, *C*_*x*_ the capacitance, 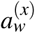 the strength of subthreshold adaptation, 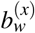 the strength of spike-triggered adaptation, 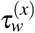 the time scale of the recovery variable and 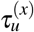 the time scale of the membrane potential (see Sl Computational Methods for parameter values). An additional term controlled by *g*_*x*_ (Eq. 1) models the regenerative activity in the dendrites as described below, but is absent from inhibitory cells (*g*_a_ = *g*_b_ = 0). Also, an additional term controlled by *c*_*x*_ and the kernel *K* (see SI Computational Methods) models how the last action potential at 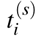 is back propagating from the soma in the dendrites, and is absent from all other units (*c*_s_,*c*_a_,*c*_b_ = 0). When units s, a and b reach a threshold at *V*_*T*_ the membrane potential is reset to the reversal potential after an absolute refractory period of 3 ms and a spike is added to the spike train 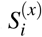 in the form of a sum of Dirac functions.

The dendritic compartment has nonlinear dynamics dictated by the sigmoidal function *f* to model the nonlinear activation of calcium channels^43^. This current propagates to the somatic unit such that *g*_s_ controls the somatic effect of forward calcium spike propagation. In the dendritic compartment, the parameter *g*_d_ controls the potency of local regenerative activity. The dendritic recovery variable controls both the duration of the calcium spike consistent with potassium currents^96,97^ and resonating subthreshold dynamics consistent with h-current^55^. Note that the differential equations are similar to those used to model NMDA spikes^98^, but the strong and relatively fast recovery variable ensures that the calcium spikes have shorter durations (10-40 ms). Membrane time constants were matched to values found in vivo^80^ (see SI Computational Methods for parameter values).

Each unit receives a combination of synaptic input, external input and background noise (see SI Computational Methods). Synapses were modeled as exponentially decaying changes in conductance. Connection probability was chosen to 0.2 for excitatory connections and one for inhibitory connections consistent with experimental observations^80,86^. Background noise was modeled as a time-dependent Ornstein-Uhlenbeck process independently drawn for each unit. We chose the amplitude of background fluctuations to be such that the standard deviation of membrane potential fluctuations is around 6 mV as observed in V1 L2-3 pyramidal neurons^46^. To model STP, we used the extended Tsodyks-Markram model^59,61^ with parameters consistent with experimental calibrations *in vitro*^61^ (see SI Computational Methods).

## Acknowledgments

We thank Guillaume Hennequin, Jean-Claude Béïque and Matthew Larkum for helpful discussions. We thank Loreen Hertäg, Alexandre Payeur and Stephen E. Clarke for critical reading of the manuscript as well as Greg Knoll for an independent verification of the numerical results. This work was supported by a Bernstein Award (01GQ1201) by the German Federal Ministry for Science and Education and an NSERC Discovery Grant 06872. Part of this work was conducted (RN and HS) at the Computational and Biological Learning Laboratory, Department of Engineering, University of Cambridge, UK.

## Author Declarations

The authors declare no conflict of interest.

## Author Contributions

HS and RN designed the study and wrote the manuscript. RN performed the simulations analyzed the results and carried out the theoretical analysis.

## References

1. Hubel, D. & Wiesel, T. Receptive fields, binocular interaction and functional architecture in the cat’s visual cortex. J Physiol 160, 106–154 (1962).

2. Felleman, D. & van Essen, D. Distributed hierarchical processing in the primate cerebral cortex. Cereb. Cortex 1, 1–47 (1991).

3. Rauschecker, J. P. & Scott, S. K. Maps and streams in the auditory cortex: nonhuman primates illuminate human speech processing. Nat. Neurosci. 12, 718–724 (2009).

4. Graziano, M. The organization of behavioral repertoire in motor cortex. Annu. Rev. Neurosci. 29, 105–134 (2006).

5. Mineault, P. J., Khawaja, F. A., Butts, D. A. & Pack, C. C. Hierarchical processing of complex motion along the primate dorsal visual pathway. Proc. Natl. Acad. Sci. USA 109, E972–E980 (2012).

6. De Pasquale, R. & Sherman, S. M. Synaptic properties of corticocortical connections between the primary and secondary visual cortical areas in the mouse. J. Neurosci. 31, 16494–16506 (2011).

7. Niell, C. M. & Stryker, M. P. Modulation of visual responses by behavioral state in mouse visual cortex. Neuron 65, 472–479 (2010).

8. Keller, G. B., Bonhoeffer, T. & Hübener, M. Sensorimotor mismatch signals in primary visual cortex of the behaving mouse. Neuron 74, 809–815 (2012).

9. Gilbert, C. D. & Li, W. Top-down influences on visual processing. Nat. Rev. Neurosci. 14, 350–363 (2013).

10. Dayan, P., Hinton, G. E., Neal, R. M. & Zemel, R. S. The helmholtz machine. Neural Comput. 7, 889–904 (1995).

11. Bastos, A. M. et al. Canonical microcircuits for predictive coding. Neuron 76, 695–711 (2012).

12. Rumelhart, D. E., Hinton, G. E. & Williams, R. J. Learning representations by back-popagating errors. Nature 323, 533–536 (1986).

13. Lillicrap, T. P., Cownden, D., Tweed, D. B. & Akerman, C. J. Random synaptic feedback weights support error backpropagation for deep learning. Nature Commun. 7 (2016).

14. Guergiuev, J., Lillicrap, T. P. & Richards, B. A. Towards deep learning with segregated dendrites. eLife (2017).

15. Crick, F. Function of the thalamic reticular complex: the searchlight hypothesis. Proc. Natl. Acad. Sci. USA 81, 4586–4590 (1984).

16. Engel, A. K. & Singer, W. Temporal binding and the neural correlates of sensory awareness. Trends Cogn. Sci. 5, 16–25 (2001).

17. Larkum, M. A cellular mechanism for cortical associations: an organizing principle for the cerebral cortex. Trends Neurosci. 36, 141–151 (2013).

18. Kayser, C., Montemurro, M., Logothetis, N. & Panzeri, S. Spike-phase coding boosts and stabilizes information carried by spatial and temporal spike patterns. Neuron 61, 597–608 (2009).

19. Akam, T. & Kullmann, D. M. Oscillatory multiplexing of population codes for selective communication in the mammalian brain. Nat. Rev. Neurosci. 15, 111 (2014).

20. O’Reilly, R. C. Biologically plausible error-driven learning using local activation differences: The generalized recirculation algorithm. Neural Computation 8, 895–938 (1996).

21. Kaifosh, P. & Losonczy, A. Mnemonic functions for nonlinear dendritic integration in hippocampal pyramidal circuits. Neuron 90, 622–634 (2016).

22. Knight, B. W. Dynamics of encoding in a population of neurons. J. Gen. Physiology 59, 734–766 (1972).

23. Fairhall, A. L., Lewen, G., Bialek, W. & van Steveninck, R. Efficiency and ambiguity in an adaptive neural code. Nature 412, 787–792 (2001).

24. Gerstner, W., Kistler, W., Naud, R. & Paninski, L. Neuronal Dynamics (Cambridge University Press, Cambridge, 2014).

25. Wang, X. et al. Feedforward excitation and inhibition evoke dual modes of firing in the cat’s visual thalamus during naturalistic viewing. Neuron 55, 465–478 (2007).

26. Kepecs, A., van Rossum, M., Song, S. & Tegner, J. Spike-timing-dependent plasticity: Common themes and divergent vistas. Biol. Cybern. 87, 446–458 (2002).

27. Larkum, M., Zhu, J. & Sakmann, B. A new cellular mechanism for coupling inputs arriving at different cortical layers. Nature 398, 338–341 (1999).

28. Shai, A. S., Anastassiou, C. A., Larkum, M. E. & Koch, C. Physiology of layer 5 pyramidal neurons in mouse primary visual cortex: coincidence detection through bursting. PLoS Comput Biol 11, e1004090 (2015).

29. Gabbiani, F., Metzner, W., Wessel, R., Koch, C. et al. From stimulus encoding to feature extraction in weakly electric fish. Nature 384, 564–567 (1996).

30. Bastian, J. & Nguyenkim, J. Dendritic modulation of burst-like firing in sensory neurons. Journal of Neurophysiology 85, 10–22 (2001).

31. Oswald, A., Chacron, M., Doiron, B., Bastian, J. & Maler, L. Parallel processing of sensory input by bursts and isolated spikes. J. Neurosci. 24, 4351–4362 (2004).

32. Deschênes, M., Paradis, M., Roy, J. & Steriade, M. Electrophysiology of neurons of lateral thalamic nuclei in cat: resting properties and burst discharges. J. Neurophys. 51, 1196–1219 (1984).

33. Brumberg, J. C., Nowak, L. G. & McCormick, D. A. Ionic mechanisms underlying repetitive high-frequency burst firing in supragranular cortical neurons. J. Neurosci. 20, 4829–4843 (2000).

34. Magee, J. C. & Carruth, M. Dendritic voltage-gated ion channels regulate the action potential firing mode of hippocampal ca1 pyramidal neurons. J. Neurophys. 82, 1895–1901 (1999).

35. Bittner, K. C. et al. Conjunctive input processing drives feature selectivity in hippocampal ca1 neurons. Nat. Neurosci. 18, 1133–1142 (2015).

36. Doiron, B., Longtin, A., Berman, N. & Maler, L. Subtractive and divisive inhibition: effect of voltage-dependent inhibitory conductances and noise. Neural Comput. 13, 227–248 (2001).

37. Körding, K. P. & König, P. Supervised and unsupervised learning with two sites of synaptic integration. J Comput. Neurosci. 11, 207–215 (2001).

38. Izhikevich, E. M. Dynamical systems in neuroscience: the geometry of excitability and bursting (MIT Press, Cambridge, Mass., 2007).

39. Giugliano, M., La Camera, G., Fusi, S. & Senn, W. The response of cortical neurons to in vivo-like input current: theory and experiment: Ii. time-varying and spatially distributed inputs. Biol. Cybern. 99, 303–318 (2008).

40. Harris, K. D., Hirase, H., Leinekugel, X., Henze, D. A. & Buzsáki, G. Temporal interaction between single spikes and complex spike bursts in hippocampal pyramidal cells. Neuron 32, 141–149 (2001).

41. Clarke, S. E. & Maler, L. Feedback synthesizes neural codes for motion. Curr. Biol. 27, 1356–1361 (2017).

42. Larkum, M. E., Kaiser, K. & Sakmann, B. Calcium electrogenesis in distal apical dendrites of layer 5 pyramidal cells at a critical frequency of back-propagating action potentials. Proc. Natl. Acad. Sci. USA 96, 14600–14604 (1999).

43. Naud, R., Bathellier, B. & Gerstner, W. Spike-timing prediction in cortical neurons with active dendrites. Front. Comput. Neurosci. 8, 90 (2014).

44. De Kock, C. & Sakmann, B. High frequency action potential bursts (> 100 hz) in l2/3 and l5b thick tufted neurons in anaesthetized and awake rat primary somatosensory cortex. J. Physiol. 586, 3353–3364 (2008).

45. Renart, A. et al. The asynchronous state in cortical circuits. Science 327, 587–90 (2010).

46. Polack, P.-O., Friedman, J. & Golshani, P. Cellular mechanisms of brain state-dependent gain modulation in visual cortex. Nat. Neurosci. 16, 1331–1339 (2013).

47. Larkum, M. E., Senn, W. & Luscher, H.-R. Top-down dendritic input increases the gain of layer 5 pyramidal neurons. Cereb. Cortex 14, 1059–1070 (2004).

48. Hay, E. & Segev, I. Dendritic excitability and gain control in recurrent cortical microcircuits. Cereb. Cortex bhu200 (2014).

49. Thorpe, S., Fize, D. & Marlot, C. Speed of processing in the human visual system. Nature 381, 520–522 (1996).

50. Bialek, W., Rieke, F., de Ruyter Van Steveninck, R. & Warland, D. Reading a neural code. Science 252, 1854–1857 (1991).

51. Pillow, J. et al. Spatio-temporal correlations and visual signalling in a complete neuronal population. Nature 454, 995–999 (2008).

52. Naud, R. & Gerstner, W. Coding and decoding with adapting neurons: A population approach to the peri-stimulus time histogram. PLOS Comput. Biol. 8, e1002711 (2012).

53. Middleton, J. W., Yu, N., Longtin, A. & Maler, L. Routing the flow of sensory signals using plastic responses to bursts and isolated spikes: experiment and theory. J. Neurosci. 31, 2461–2473 (2011).

54. Deger, M., Schwalger, T., Naud, R. & Gerstner, W. Fluctuations and information filtering in coupled populations of spiking neurons with adaptation. Phys. Rev. E 90, 062704 (2014).

55. Kole, M. H. P., Hallermann, S. & Stuart, G. J. Single ih channels in pyramidal neuron dendrites: properties, distribution, and impact on action potential output. J. Neurosci. 26, 1677–87 (2006).

56. Wei, W. & Wolf, F. Spike onset dynamics and response speed in neuronal populations. Phys. Rev. Lett. 106, 088102 (2011).

57. Pérez-Garci, E., Gassmann, M., Bettler, B. & Larkum, M. The gabab1b isoform mediates long-lasting inhibition of dendritic ca2+ spikes in layer 5 somatosensory pyramidal neurons. Neuron 50, 603–616 (2006).

58. Markram, H., Wu, Y. & Tosdyks, M. Differential signaling via the same axon of neocortical pyramidal neurons. Proc. Natl. Acad. Sci. USA 95, 5323–5328 (1998).

59. Markram, H., Wang, Y. & Tsodyks, M. Differential signaling via the same axon of neocortical pyramidal neurons. Proc. Nal. Acad. Sci. USA 95, 5323–5328 (1998).

60. Izhikevich, E. M., Desai, N. S., Walcott, E. C. & Hoppensteadt, F. C. Bursts as a unit of neural information: selective communication via resonance. Trends Neurosci. 26, 161–167 (2003).

61. Costa, R. P., Sjöström, P. J. & Van Rossum, M. C. Probabilistic inference of short-term synaptic plasticity in neocortical microcircuits. Front. Comput. Neurosci. 7 (2013).

62. Silberberg, G. & Markram, H. Disynaptic inhibition between neocortical pyramidal cells mediated by martinotti cells. Neuron 53, 735–746 (2007).

63. Murphy, B. K. & Miller, K. D. Multiplicative gain changes are induced by excitation or inhibition alone. J. Neurosci. 23, 10040–10051 (2003).

64. Murayama, M. et al. Dendritic encoding of sensory stimuli controlled by deep cortical interneurons. Nature 457, 1137–1141 (2009).

65. Gidon, A. & Segev, I. Principles governing the operation of synaptic inhibition in dendrites. Neuron 75, 330–341 (2012).

66. Royer, S. et al. Control of timing, rate and bursts of hippocampal place cells by dendritic and somatic inhibition. Nat. Neurosci. 15, 769–775 (2012).

67. Xie, X. & Seung, H. S. Equivalence of backpropagation and contrastive hebbian learning in a layered network. Neural Comput. 15, 441–454 (2003).

68. Womelsdorf, T., Ardid, S., Everling, S. & Valiante, T. A. Burst firing synchronizes prefrontal and anterior cingulate cortex during attentional control. Current Biology 24, 2613–2621 (2014).

69. Tian, L. et al. Imaging neural activity in worms, flies and mice with improved gcamp calcium indicators. Nature Methods 6, 875–881 (2009).

70. Mease, R. A., Kuner, T., Fairhall, A. L. & Groh, A. Multiplexed spike coding and adaptation in the thalamus. Cell Reports 19, 1130–1140 (2017).

71. Larkum, M., Nevian, T., Sandler, M., Polsky, A. & Schiller, J. Synaptic integration in tuft dendrites of layer 5 pyramidal neurons: a new unifying principle. Science (2009).

72. Gerstner, W. Population dynamics of spiking neurons: Fast transients, asynchronous states, and locking. Neural Comput. 12, 43–89 (2000).

73. Sompolinsky, H., Yoon, H., Kang, K. & Shamir, M. Population coding in neuronal systems with correlated noise. Phys. Rev. E 64, 051904 (2001).

74. Cannon, R. C., O’Donnell, C. & Nolan, M. F. Stochastic ion channel gating in dendritic neurons: morphology dependence and probabilistic synaptic activation of dendritic spikes. PLoS Comput. Biol. 6, e1000886 (2010).

75. O’Donnell, C. & van Rossum, M. C. Systematic analysis of the contributions of stochastic voltage gated channels to neuronal noise. Front. Comp. Neurosci. 8, 105 (2014).

76. Naud, R., Payeur, A. & Longtin, A. Noise gated by dendrosomatic interactions increases information transmission. Phys. Rev. X 7, 031045 (2017).

77. van Vreeswijk, C. & Sompolinsky, H. Chaos in neuronal networks with balanced excitatory and inhibitory activity. Science 274, 1724–1726 (1996).

78. Vogels, T. P., Sprekeler, H., Zenke, F., Clopath, C. & Gerstner, W. Inhibitory plasticity balances excitation and inhibition in sensory pathways and memory networks. Science 334, 1569–1573 (2011).

79. Reyes, A. et al. Target-cell-specific facilitation and depression in neocortical circuits. Nat. Neurosci. 1, 279–285 (1998).

80. Pala, A. & Petersen, C. C. In vivo measurement of cell-type-specific synaptic connectivity and synaptic transmission in layer 2/3 mouse barrel cortex. Neuron 85, 68–75 (2015).

81. Lee, S., Kruglikov, I., Huang, Z. J., Fishell, G. & Rudy, B. A disinhibitory circuit mediates motor integration in the somatosensory cortex. Nat. Neurosci. 16, 1662–1670 (2013).

82. Pi, H.-J. et al. Cortical interneurons that specialize in disinhibitory control. Nature 503, 521–524 (2013).

83. Pfeffer, C. K., Xue, M., He, M., Huang, Z. J. & Scanziani, M. Inhibition of inhibition in visual cortex: the logic of connections between molecularly distinct interneurons. Nat. Neurosci. 16, 1068–1076 (2013).

84. Jiang, X. et al. Principles of connectivity among morphologically defined cell types in adult neocortex. Science 350, aac9462 (2015).

85. Markram, H. et al. Interneurons of the neocortical inhibitory system. Nat Rev Neurosci 5, 793–807 (2004).

86. Fino, E. & Yuste, R. Dense inhibitory connectivity in neocortex. Neuron 69, 1188–1203 (2011).

87. Adesnik, H., Bruns, W., Taniguchi, H., Huang, Z. J. & Scanziani, M. A neural circuit for spatial summation in visual cortex. Nature 490, 226–231 (2012).

88. Cruikshank, S. J., Urabe, H., Nurmikko, A. V. & Connors, B. W. Pathway-specific feedforward circuits between thalamus and neocortex revealed by selective optical stimulation of axons. Neuron 65, 230–245 (2010).

89. Rojas-Piloni, G. et al. Relationships between structure, in vivo function and long-range axonal target of cortical pyramidal tract neurons. Nature Commun. 8, 1–11 (2017).

90. Petreanu, L., Mao, T., Sternson, S. M. & Svoboda, K. The subcellular organization of neocortical excitatory connections. Nature 457, 1142–5 (2009).

91. Xu, H., Jeong, H.-Y., Tremblay, R. & Rudy, B. Neocortical somatostatin-expressing gabaergic interneurons disinhibit the thalamorecipient layer 4. Neuron 77, 155–167 (2013).

92. Van Kerkoerle, T., Self, M. W. & Roelfsema, P. R. Layer-specificity in the effects of attention and working memory on activity in primary visual cortex. Nat. Commun. 8, 13804 (2017).

93. Takahashi, N., Oertner, T. G., Hegemann, P. & Larkum, M. E. Active cortical dendrites modulate perception. Science 354, 1587–1590 (2016).

94. Wörgötter, F., Nelle, E., Li, B. & Funke, K. The influence of corticofugal feedback on the temporal structure of visual responses of cat thalamic relay cells. J. Physiol. 509, 797–815 (1998).

95. Ortuño, T., Grieve, K. L., Cao, R., Cudeiro, J. & Rivadulla, C. Bursting thalamic responses in awake monkey contribute to visual detection and are modulated by corticofugal feedback. Front. Behav. Neurosci. 8 (2014).

96. Cai, X. et al. Unique roles of sk and kv4. 2 potassium channels in dendritic integration. Neuron 44, 351–364 (2004).

97. Harnett, M. T., Xu, N.-L., Magee, J. C. & Williams, S. R. Potassium channels control the interaction between active dendritic integration compartments in layer 5 cortical pyramidal neurons. Neuron 79, 516–529 (2013).

98. Major, G., Polsky, A., Denk, W., Schiller, J. & Tank, D. W. Spatiotemporally graded nmda spike/plateau potentials in basal dendrites of neocortical pyramidal neurons. J Neurophysiol 99, 2584–601 (2008).

